# Impact of α-synuclein fibrillar strains and ß-amyloid assemblies on the endolysosomal logistics of mouse cortical neurons

**DOI:** 10.1101/2021.05.15.444288

**Authors:** Qiao-Ling Chou, Ania Alik, François Marquier, Ronald Melki, François Treussart, Michel Simonneau

## Abstract

Endosomal transport and positioning are involved in establishing neuronal compartment architecture, dynamics and function, contributing to neuronal intracellular logistics. Furthermore, endo-lysosomal dysfunction has been identified as a common mechanism in neurodegenerative diseases. Here, we analyzed endolysosomal transport following the external application of α-synuclein (α-syn) fibrillar polymorphs, ß-amyloid (Aß) fibrils and oligomers on primary cultures of mouse cortical neurons. We used a simple readout to measure this transport: the spontaneous endocytosis of fluorescent nanodiamonds — a perfectly stable nano-emitter — in cultured neurons. We then performed a high-throughput automatic extraction and quantification of the directed motions of these nanodiamonds. α-syn fibrillar polymorphs, Aß fibrils and oligomers halved the proportion of nanodiamonds transported along microtubules, but only slightly decreased their interactions with cortical neurons. This large decrease in endosomal transport would be expected to have a huge impact on neuronal homeostasis. We then assessed lysosomal dynamics with Lysotracker. The exposure of neurons to Aß oligomers led to an increase in the number of lysosomes, a decrease in the fraction of moving lysosomes and an increase in their size, reminiscent of findings for the APP transgenic model of Alzheimer’s disease. We then analyzed the effect of α-syn fibrillar polymorphs, Aß fibrils and oligomers on endosomal and lysosomal transport and quantified the directed transport of these assemblies within cortical neurons. We report different impacts on endosomal and lysosomal transport parameters and differences in trajectory length for cargoes loaded with pathogenic protein assemblies. Our results suggest that the internalization and transport of intraneuronal pathogenic protein aggregates are potential targets for novel neuroprotective treatment strategies.

**Significance Statement:** Neurodegenerative diseases (NDs) are characterized by the deposition of protein aggregates with broad-range neuronal toxicity. Defects of endolysosomal trafficking are increasingly being seen as key pathological features of NDs, probably contributing to synaptic dysfunction and ultimate neuronal death. We used fast fluorescence videomicroscopy to investigate endosomal and lysosomal dynamics in the branches of mouse cortical neurons in primary cultures following the application of α-syn fibrillar polymorphs (fibrils and ribbons) and Aß assemblies (oligomers and fibrils). We provide new insight into the differential effects of these pathogenic protein assemblies on endosomal and lysosomal transport, and reveal differences in the transport characteristics of the compartments loaded with these protein assemblies relative to endosomes.

## Introduction

The impairment of axonal transport has recently emerged as a factor common to several neurodegenerative disorders (Millecamps and Julien, 2013; Morfini et al., 2009). This has led to suggestions that an early impact on intraneuronal transport may be a phenotypic trait common to neurodegenerative diseases such as Alzheimer’s, Huntington’s and Parkinson’s diseases (Stokin et al., 2005; Saudou and Humbert, 2016; Volpicelli-Daley et al., 2014). There is compelling evidence that abnormal protein accumulation in the brain is a key pathophysiological mechanism underlying the neurotoxicity observed in these age-related disorders (Golde et al., 203; Soto et al., 2018). The selective aggregation of misfolded proteins is a hallmark of these neurodegenerative diseases (Saez-Atienzar & Masliah, 2020). However, different species of the same molecule, such as oligomers and fibrils, may contribute to a whole spectrum of toxicities, adding a level of complexity (Alam et al., 2017).

A few studies have compared fibrillary and oligomeric α-synuclein (α-syn) and ß-amyloid (Aß) — known to be involved in Parkinson’s disease and Alzheimer’s disease, respectively — within the same neurons. For example, Brahic et al. (2016) demonstrated the anterograde and retrograde transport of α-syn, Aß_42_ and HTTExon1 fibrils with various efficiencies in the axons of mouse primary neurons grown in microfluidic chambers. Here, we quantified, in detail, the impact of two α-syn fibrillar polymorphs — fibrils (α-synF) and ribbons (α-synR) — Aß_42_ fibrils (AßF) and oligomers (AßO) on endosomal and lysosomal transport in primary cultures of mouse neurons. We used a method established in a previous study to measure this transport and investigate its parameters (Haziza et al., 2017). This method involved the spontaneous internalization, by neurons, of stable, non-toxic, fluorescent nanodiamonds (FNDs) in endosomes. The displacement of the nanodiamonds was followed by fast video microscopy, and single-particle trajectories were then automatically extracted from the videos and analyzed. We then used fluorescently labeled α-syn and Aß assemblies to perform similar investigations to investigate the intraneuronal transport of these assemblies.

Our data make it possible to address two complementary questions: (i) Do α-synF, α-synR, AßF and AßO influence the fraction of cargoes moving along the microtubules? (ii) Do they affect the dynamics of intracellular endosomal and lysosomal transport? We show here that all the pathogenic protein assemblies tested decrease the fraction of endosomes moving along microtubules and affect some of their transport parameters. In addition, AßO also affected the properties of lysosomes (number, fraction of lysosomes transported, and transport parameters). Finally, our data indicate that cargoes loaded with α-synF, α-synR, AßF or AßO are transported differently from endosomes (control cargoes). This suggests that cargo-motor assemblies may have different molecular characteristics.

## Material and Methods

### Production of α-syn fibrillar assemblies, Aß fibrils and oligomers

The expression and purification of human WT α-syn was performed as previously described (Ghee et al., 2005). Pure WT α-syn was incubated in buffer A to obtain the fibrillar polymorph ‘‘fibrils’’ α-synF (50 mM Tris-HCl at pH 7.5, 150 mM KCl) and in buffer B for ‘‘ribbons’’ α-synR (5 mM Tris-HCl at pH 7.5) at 37°C under continuous shaking in an Eppendorf Thermomixer (Hamburg, Germany) set at 600 rotations per minute (rpm) for 4–7 days (Bousset et al., 2013). The fibrillar α-syn polymorphs were centrifuged twice at 15,000 *g* for 10 min and resuspended twice in phosphate-buffered saline (PBS) at 1,446 g/L prior to labeling with ATTO 488 NHS-ester (#AD 488-3, Atto-Tec, Siegen, Germany) fluorophore following the manufacturer’s instructions using a protein/dye ratio of 1:2. The labeling reactions were arrested by addition of 1 mM Tris (pH 7.5). The unreacted fluorophore was removed by a final cycle of two centrifugations at 15,000 *g* for 10 min and resuspensions of the pellets in PBS. This labeling protocol typically yields ≥ 1 ATTO molecule incorporated per α-syn monomer on average as previously demonstrated (Shrivastava et al., 2015). The assemblies were examined by transmission electron microscopy after adsorption on 200 mesh carbon-coated electron microscopy grids and negative stained with 1% uranyl acetate before and after fragmentation using a JEOL 1400 electron microscope (JEOL, Tokyo, Japan).

The expression and purification of Met-Aß 1-42 was performed as described (Walsh et al., 2009). Aß was assembled in PBS, at 4°C or 37°C without shaking for 2 or 24 h to obtain oligomers AßO or fibrils AßF, respectively. The two kinds of assemblies were labeled with ATTO 488 NHS-ester at a protein/dye ratio of 1:2. The labeling reactions were arrested by addition of 1 mM Tris at pH 7.5. For fibrillar Aß, the unreacted fluorophore was removed by two cycles of centrifugation and resuspension of the pelleted fibrils in PBS as described for α-syn. For oligomeric Aß, the oligomers were separated from the monomeric and fibrillar forms of the protein by size exclusion chromatography on a Superose 6 HR10/300 column (GE Healthcare, Life Sciences, Wauwatosa, WI, USA) equilibrated in PBS pH 7.4 at a flow rate of 0.5 mL/min. Elution was monitored by measuring absorbance at 280 nm wavelength. The Superose 6 column was calibrated with Dextran blue (over 2200 kDa), (670 kDa), β -amylase (200 kDa), BSA (66 kDa), and carbonic anhydrase (29 kDa) standards (Sigma-Aldrich).

### Primary mouse cortical neuron cultures

We used commercial primary mouse cortical neurons (ref. Gibco A15586, ThermoFisher) because the provider quality check guarantees a purity of 98% of neurons. The cells were grown on high optical quality glass coverslips (high-precision 170±5 μm thick, 18 mm diameter, ref. 0117580, Marienfeld GmbH, Germany). The coverslips are first cleaned with 70% ethanol, rinsed with water for injection (ref. A128730, ThermoFisher Inc., USA) and exposed during 1 h to UV light. They were then coated with 0.1 mg/ml poly-L-ornithine (ref. P3655, Sigma-Aldrich Merck KGaA, Germany) and placed for 2 h in an incubator set at 37°C, then rinsed twice with water and let dry at biological hood for one hour. We plated an amount of 6×10^5^ primary mouse cortical neurons (ref. Gibco A15586, ThermoFisher) on each coated coverslip, which was then put at the bottom of a 6-wells plate, each well-being finally filled with 3 mL of neurobasal phenol red-free medium (ref. 12348017, ThermoFisher) containing 0.5 mM GlutaMax (ref. 35050061 ThermoFisher), 2% B-27 (ref. 17504044, ThermoFisher) and 1% PenStrep (ref. 15070063, ThermoFisher). The 6-well plate was then placed in an incubator at 37 °C and 5% CO_2_. Half of the volume of the medium was replaced with fresh medium 24 h after plating. We made the subsequent medium changes every 3 days to reduce glutamate toxicity. Neurons were grown until 21 days in culture.

### Exposure of mouse cortical neurons to α-syn fibrillar assemblies, Aß fibrils or oligomers

In all the measurements dealing with (1) the impact of pathogenic protein assemblies on endosomal-lysosomal transport, (2) their colocalization with FND-labelled compartment or lysosomes, or (3) the tracking of their intraneuronal transport by fluorescence videomicroscopy, cortical neurons were incubated with either 0.2 μM ATTO 488-labeled α-synF or R, or 1 μM ATTO 488-labeled AßF or AßO. α-synF or R were added at 24 h, 48 h, or 72 h before observations, while the addition time was either 24 h or 48 h for ATTO 488-labeled AßF or AßO. The video acquisitions of all the experiments were performed at DIC21.

### Washing protocol to test the protein assemblies interaction with the neuron membrane

The coverslip with the culture attached to them, were extracted from the well and flushed with PBS first and then twice with culture medium, before FND internalization was carried out following the procedure described in the next paragraph.

### Intraneuronal transport cargo labeling

To evaluate the endosomal transport parameters, we relied on our fluorescent nanodiamond assay (Haziza *et al*. 2017). We used commercially available sized 35 nm FND (brFND-35, FND Biotech, Taiwan). Each NP contains an average of 15 nitrogen-vacancy emitters displaying a peak emission wavelength around 700 nm and a full-width at half-maximum of ≈100 nm. This far-red emission allows also to investigate the colocalization of green-emitting ATTO 488-labelled neurodegenerative-disease related species with FND-labeled cargos. FND were internalized in cortical neurons just before the transport analysis, at DIC21. Each culture coverslip was removed from the 6-well plate containing maintaining medium and put in contact with 400 μL of fresh culture medium to which we added 2 μl of stock solution of FNDs (1 mg/mL), reaching a final FND concentration of 5 μg/mL. After 10 mins incubation, the extra FND-containing medium was absorbed by a wiper sheet and the coverslip was placed back to the dish containing the old maintaining medium. The culture was then placed back during 20 mins in the incubator before the video acquisition.

To measure lysosomal transport or investigate the colocalization of neurodegenerative disease-related species with lysosomes, cortical neurons were stained at DIC21, just before the observation, with LysoTracker Deep Red (ref. L12492, ThermoFisher) or Magic Red Cathepsin B substrate (ref. ICT937, Bio-Rad). These dye molecules have an emission spectrum within the similar range than the one of FND. The coverslip was removed from maintaining medium and incubated with prewarmed (37°C) culture medium containing 50 nM LysoTracker or Magic Red (1:20 dilution) for 1 h. The probe-containing medium was replaced with the old maintaining medium and followed by video acquisition.

### Pseudo-total internal reflection (TIRF) live-cell videomicroscopy

Pseudo-TIRF illumination was implemented on an inverted microscope (Eclipse Ti-E, Nikon, Japan) as described in details in (Haziza et al. 2017). The whole microscope is enclosed in a cage incubator (Okolab, Italy) to maintain temperature at 37°C. For the intraneuronal transport recording, each coverslip supporting the neuron culture is mounted at the bottom a Ludin chamber (type 1, Life Imaging Service, Switzerland), installed inside the environmental chamber (in which 5% partial CO_2_ pressure and 100% hydrometry is maintained) having a hole at its bottom allowing direct optical access of the microscope objective to the coverslip. We used a ×100 magnification and 1.49 numerical aperture immersion oil objective (CFI Apo TIRF ×100 Oil, Nikon), compatible with differential inference contrast (DIC) mode. Field of views of interest of the neuron cultures were selected in white-light illumination DIC mode. Two continuous-wave lasers are coupled to the microscope and fluorescence was recorded on a cooled EMCCD array detector (DU-885K-CS0, Andor Technologies, UK) of 1004×1002 pixels, with 80 nm pixel size in the sample plane. Two-minutes duration videos were acquired at 20 full frame/s rate large enough to be able to detect short pausing duration in cargoes displacements. EMCCD parameters were selected to provide the largest signal-to-background ratio for FND label tracking at the selected frame rate, leading to EM gain of 90, preamplification gain ×3.8, and digitalization speed of 35 MHz. FNDs and Lysotracker fluorophore were excited with a diode-pumped solid-state laser at a wavelength of 561 nm (SLIM-561-100, Oxxius S.A., France), while ATTO 488 dye was excited with a laser diode emitting at a wavelength of 488 nm (LBX-488-200-CSB-PP, Oxxius S.A). Each excitation laser power was adjusted so that the detection dynamic range of all channels was identical for the above mentioned fixed EMCCD settings. This leads to 561 nm laser excitation power of 60 mW for FND, 200 μW for LysoTracker and 1 mW for Magic Red, and to 488 nm laser excitation power of 200 μW for ATTO 488 dye.

To perform two-color acquisitions and record simultaneously FND (or LysoTracker) and ATTO 488 we combined the two laser beams with a dual-band dichroic filter (ref. Di01-R488/561, Semrock, USA), and placed a dual imaging system (W-VIEW GEMINI, Hamamatsu, Japan) in front of the EMCCD array detector (DU-885K-CS0, Andor Technologies, UK). This system splits half of the detection field of view (FoV) in two color channels with a dichroic beamsplitter (FF560-FDi01, Semrock) and projects each color on half of the array detector, further preceded by bandpass detection filters (red channel: HC697/75, Semrock; green channel: ET525/50, Chroma Corporation, USA). The result is that each frame of the video contains the same rectangular FoV (1004×501 pixels) in green (ATTO 488) and red (FND and LysoTracker) emission range, allowing to identify spots that colocalize dynamically.

### Video processing and intraneuronal transport quantification

Two programs written in python were developed to extract quantitative parameters from videos automatically. The first one relies on Trackpy 0.4.2 package (Trackpy 2019), from which it uses two functions: locate to identify isolated spots in each fluorescence frame and fit them with gaussians, and link which connects the spots between frames to form trajectories using Crocker-Grier algorithm (Crocker & Grier 1996). Transport parameters are then calculated with a second program which first parses each trajectory into “go” and “stop” phases based on the confinement ratio calculation as described in (Haziza et al., 2017). Four main transport parameters are extracted for each trajectory: velocity, which is the average speed of all go phases; run length: average distance traveled during all go phases; pausing time: average duration of the stop phases, and pausing frequency (events/min). In addition to these four main parameters, we also calculated the total length of the trajectory as the sum of all run lengths during go phases.

### Lysosomes size estimation

To get an estimate of the lysosome size from the diffraction limited fluorescence images, we considered the LysoTracker spots as the result of the convolution of the microscope point spread function (assimilated to a gaussian of standard deviation, SD, *σ*_PSF_) and the lysosome assimilated to a symmetrical gaussian of SD *σ*_L_. The result of this convolution is also a gaussian of SD *σ*_T_, related to *σ*_L_ and *σ*_PSF_ by 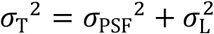. *σ*_T_ is the so-called “size” output of the Trackpy locate function. The knowledge of *σ*_T_ and *σ*_PSF_ allows the derivation of *σ*_L_. We then defined the lysosome “diameter” *d*_L_ as the full width at half maximum of its gaussian approximation, inferred by 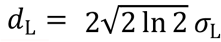. For the PSF size *σ*_PSF_ we took the measured value of the smallest FND spot observed in several trajectories with our microscope. This value was *σ*_PSF_=112 nm, consistent with the theoretical Airy radius *ρ*_A_ =286 nm (diffraction limit at 700 nm maximum emission wavelength for the 1.49 numerical aperture objective used), and the empirical relation 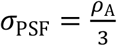, giving here an experimental PSF SD of 286/3=95 nm.

### Quantification of fluorescence intensity of ATTO 488-labeled α-synF in cortical neuron branches

We quantify the fluorescence intensity from the first frame of the green channel videos, using Fiji Image J software (Schindelin et al, 2012). We first identify from the DIC image some well-separated and mainly straight neuronal branches, that we surround with the region-of-interest (ROI) polygonal selection tool as close as possible to the branch to include all the fluorescence signal, over a length of 30 μm. We then used the Analyze function to measure the average intensity counts per pixel in the defined ROI, to which we subtract the average background counts, measured after having moved the ROI in a region without branches.

### Data representation and statistical analysis

All bar plots display the ± standard error on the mean of the distribution. Box plots display the median value as the horizontal line within the box whose limits are 25% and 75% percentiles; bottom and top horizontal lines correspond to 10% and 90% percentiles. As all the data compared between two conditions were random and normally distributed but with unequal variance (as tested with a *F*-test), we performed the relevant comparison test which is the non-parametric Wilcoxon Mann-Whitney two-tailed (implemented in Igor Pro 8, Wavemetrics Inc., USA). Stars referred to the following *p*-value significance level: \**p*<0.05; \*\**p*<0.01; \*\*\**p*<0.001.

## Results

### Quantification of intraneuronal transport with fluorescent nanodiamonds

We quantified intraneuronal transport with our FND tracking assay (Haziza et al., 2017). We first used a simple readout, by counting the FNDs detected in fields-of-view (FoV) 40×80 μm in size over a period of two minutes. The FoV selected had approximately the same density of neuronal branches, as estimated from differential interference contrast images (Fig. 1). Our incubation protocol was designed to limit any non-specific interactions of FNDs, such as attachment to the coverslip supporting the culture, and to favor the interaction of FNDs with neuron membranes and their subsequent internalization by endosomes (Haziza et al., 2017). The perfectly stable fluorescence of FNDs made it possible to reconstruct endosome trajectories accurately and to identify “stop” and “go” (no movement or very slow motion) phases (Fig. 1B) of FND-labeled endosome transport along neuronal branches by differential interference contrast microscopy (Fig. 1C).

**Figure 1.**
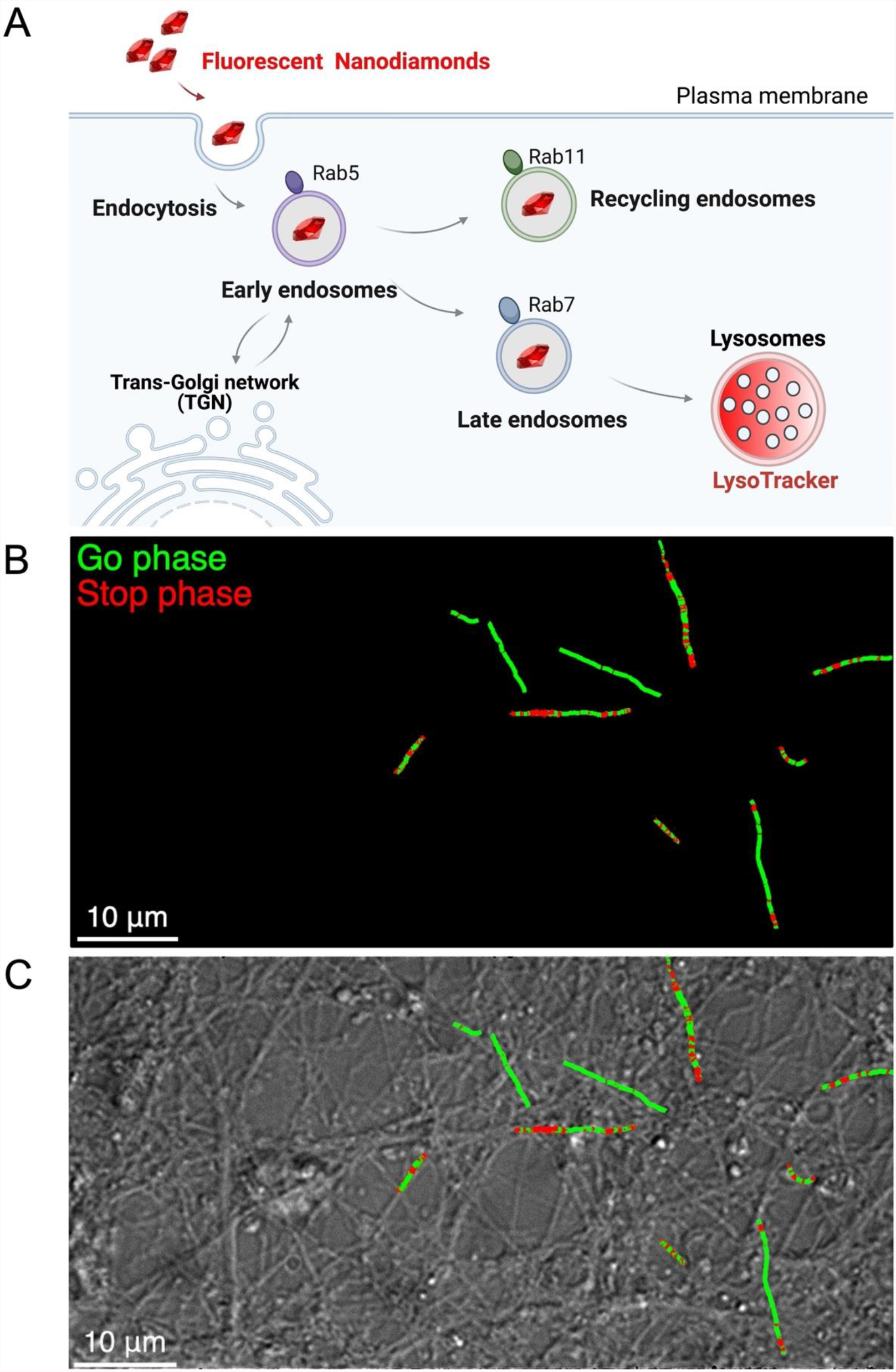
Recording fluorescent nanodiamond trajectories in mouse cortical neurons at DIC21. ***A***, Schematic representation of the cargoes that are tracked thanks to FND, at different stages after their endocytosis. We previously showed (Haziza et al., 2017) that FND are present in cargos at different stage of their lifetime after endocytosis, as shown by colocalization measurements with specific membrane protein markers: Rab5 for early endosome; Rab7 for late endosome; Rab11 for recycling endosome; Lysotracker for the lysosome. ***B***, Illustration of FND trajectories with go (green) and stop (red) phases. ***C***, Differential interference contrast images of cortical neurons overlapped with 10 representative trajectories. Scale bar: 10 μm.

### α-synF and R affect the number of cargoes transported along microtubules but do not greatly modify trajectory length

On day 20 in culture (DIC20), α-synF or R was added to primary cultures of mouse cortical neurons at a concentration of 0.2 μM, and cultures were incubated for a further 24 hours. Intraneuronal transport in these cultures was investigated on DIC21.

Exposure to either α-synF or R led to a small decrease (26% for α-synF and 13% for α-synR) in the number of FNDs (moving or not moving) present in each FoV (Fig. 2*A*). Both these fibrillar polymorphs therefore affect FND binding to neuronal membranes and FND transport dynamics within neurons. The FNDs with directed motion are those first taken up by endosomes and handled by molecular motors. This fraction of FNDs was 49% smaller after neuron exposure to α-synF, and 45% smaller after exposure to α-synR (Fig. 2*B*). The mechanism underlying this large decrease in the number of cargoes transported along microtubules is unknown, but would be expected to impair cortical neuron functions.

**Figure 2.**
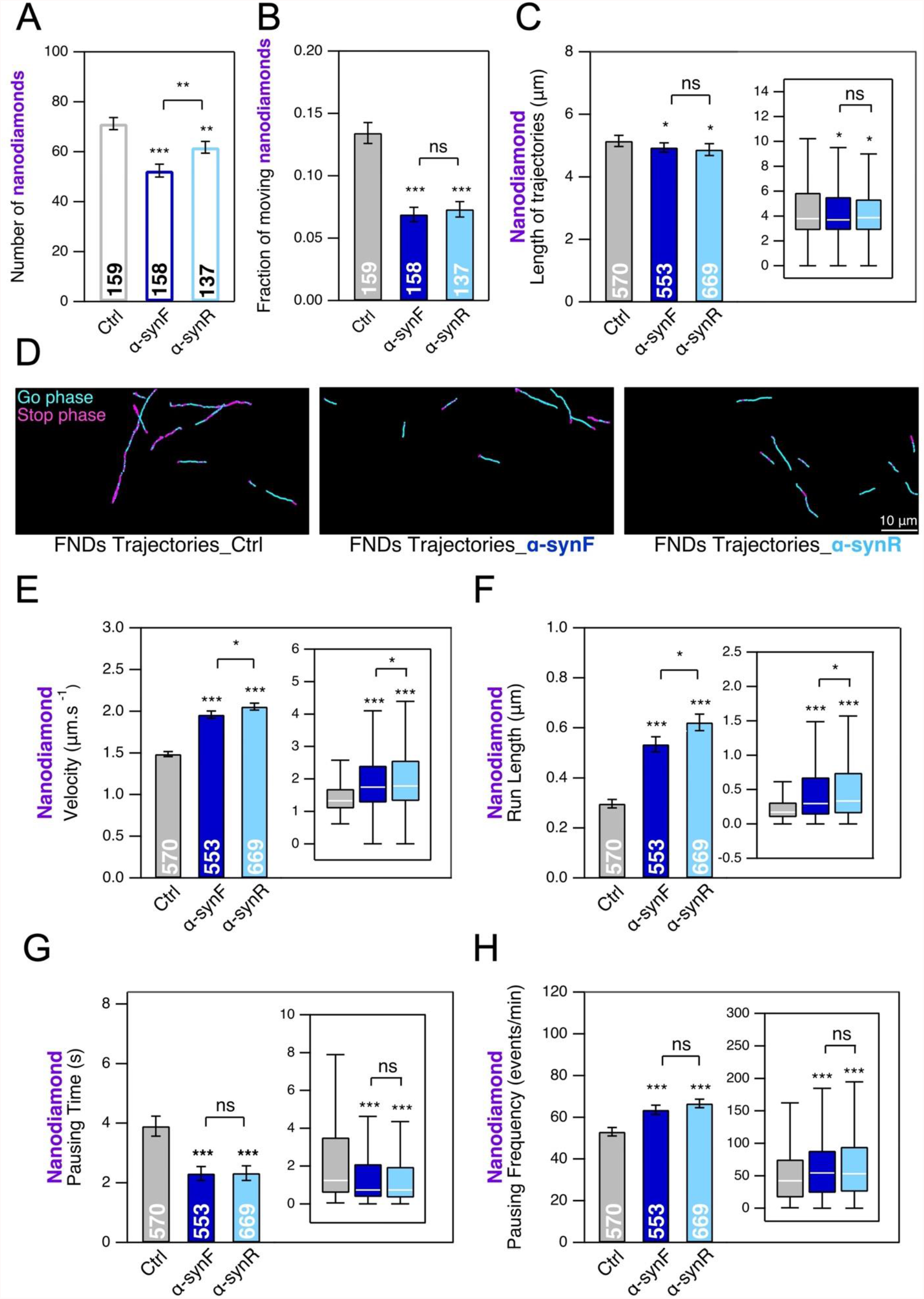
Effect of α-synF or R on the mobility of endosomes and their transport as measured by tracking FND-containing cargoes in mouse cortical neurons at DIC21. 24 h exposure to α-synF or R at 0.2 μM concentration, compared to nothing added control (Ctrl). ***A***, Number of FNDs detected per field-of-view of 40 μm x 80 μm size during 2 mins of observation. ***B***, Fraction of FNDs-containing cargoes having a directed motion. ***C***, Length of FND trajectories. ***D***, Examples of FND trajectories. Scale bar: 10 μm. ***E*-*H***, Comparison of four transport parameters: ***E***, curvilinear velocity, ***F***, run length, ***G***, pausing time and ***H***, pausing frequency. The number within each bar represents the total number of FoV (***A, B***) or trajectories (***C, E***-***H***) analyzed from *n*=8 coverslips (four independent cultures). Inset: box-plots representation of the same dataset. See also Figure 2-1 and Figure 2-2.

We investigated whether this large decrease in the moving fraction of FNDs was related to the binding of the protein assemblies to the neuronal branch membrane, possibly leading to decreased in endocytosis. For α-synF, we tried to wash off the aggregates just before adding FNDs (Materials and Methods), but we found that washing had no effect on the interaction between nanodiamonds and neurons (Fig. 2-1*A*-*C*). In particular, washing did not affect the fraction of moving FND. We concomitantly used dye fluorescence intensity methods to quantify the ATTO 488-labeled assemblies along the neuronal branches. Again, the results obtained before and after washing were not significantly different (Fig. 2-1*D*-*E*), consistent with the FND-neuron interaction results and indicating strong binding of α-synF to the neuronal membrane.

We used our established FND-based intraneuronal transport assay (Haziza et al., 2017) to detect and quantify the alternation of movement and pause phases in intraneuronal cargo transport. We first measured the length of trajectories (see Materials & Methods) for control FNDs, and for FNDs in the presence of either α-synF or α-synR. We found no major differences between the control and treated FNDs (Fig. 2*C*, Fig. 2*D*; 4% decrease for α-synF and 5% decrease for α-synR).

We then measured four parameters: the curvilinear velocity of each moving phase, its run length, the duration of pauses, and pausing frequency. Curvilinear velocity (Fig. 2*E*) and run length (Fig. 2*F*) increased (velocity: +31% and +38%; run length: +80% and +100%, for α-synF and R, respectively), whereas pausing time decreased (Fig. 2*G*; 40% decrease for both fibrillar assemblies), and pausing frequency increased (Fig. 2*H*; +19% and +25%, for α-synF and R respectively). These results are summarized in Fig. 2-2A.

We then analyzed the same parameters for lysosomes labeled with Lysotracker red, a widely used lysosome marker differing from FNDs in that all fluorescent spots, whether moving or static, correspond to lysosomes because Lysotracker is fluorescent only within lysosomes. α-synR treatment induced a slight but significant decrease (15%) in the number of lysosomes per field-of-view, whereas α-synF did not (Fig. 3*A*). This finding suggests that α-synR decreased endocytosis and is consistent with the observed decrease in FND interactions with neurons (Fig. 2*A*), possibly indicating lower levels of uptake. Furthermore, as for endosomes (Fig. 2*B*), α-synF and R induced 46% and 32% decreases, respectively, in the fraction of lysosomes displaying directed motion (Fig. 3*B*). Treatment decreased trajectory lengths slightly (Fig. 3*C*; 9% and 3% decreases for α-synF and R, respectively). Examples of FoV displaying lysosome trajectories in different conditions are shown in Fig. 3*D*, in which the large decrease in the fraction of lysosomes with directed motion is clearly visible.

**Figure 3.**
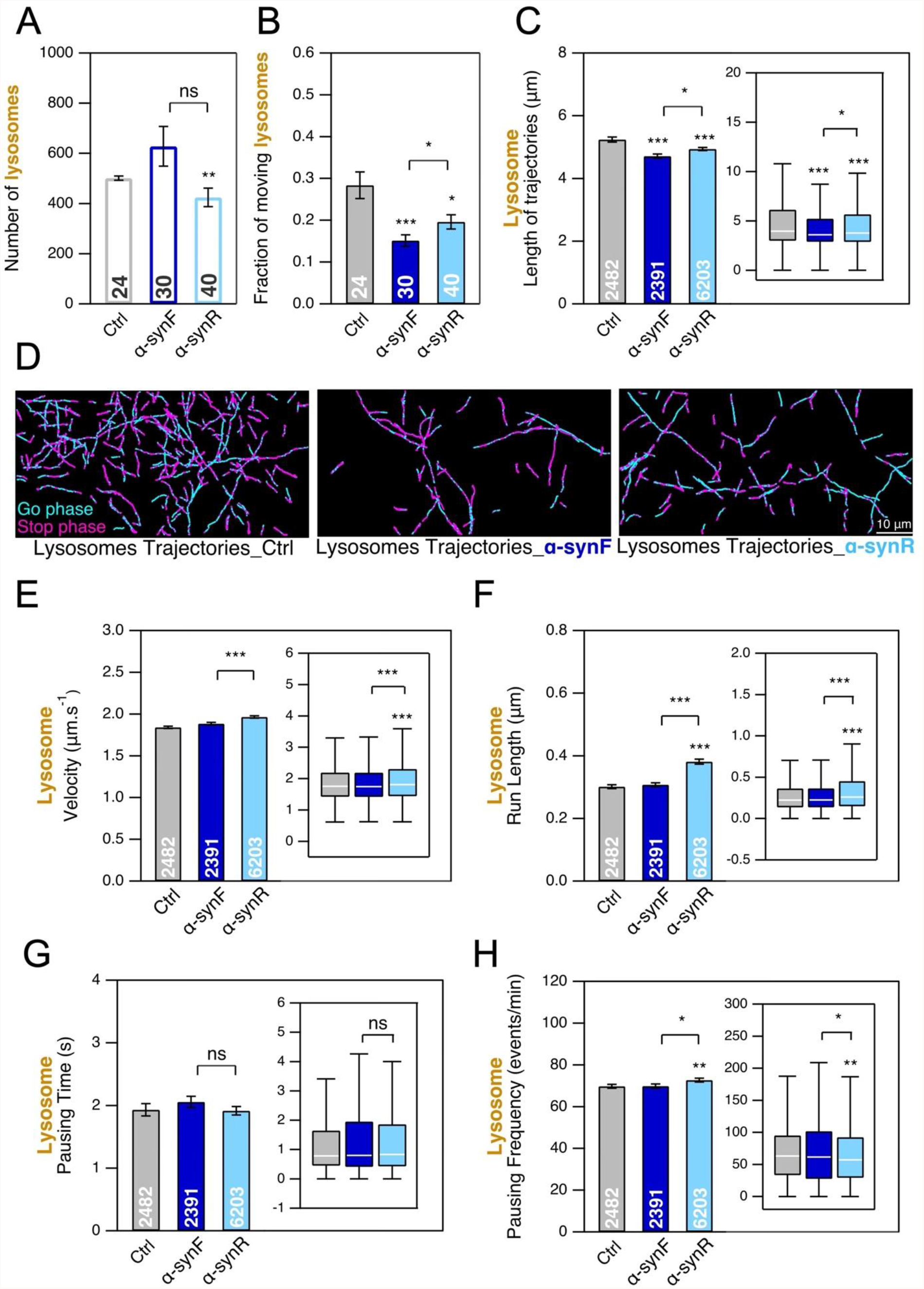
Effect of α-synF or α-synR on the mobility of LysoTracker-labelled lysosomes and their transport in mouse cortical neurons at DIC21. 24 h exposure to α-synF or α-synR at 0.2 μM concentration, compared to nothing added control (Ctrl). ***A***, Number of lysosomes detected per field-of-view of 40 × 80 μm size during 2 mins of observation. ***B***, Fraction of lysosomes having a directed motion. ***C***, Length of lysosome trajectories. ***D***, Examples of lysosome trajectories. Scale bar: 10 μm. ***E*-*H***, Comparison of four transport parameters: curvilinear velocity (***E***), run length (***F***), pausing time (***G***) and **H**, pausing frequency (**H**). The number within each bar represents the total number of FoV (***A, B***) or trajectories (***C, E***-***H***) analyzed from *n*=2 coverslips (from one culture). Inset: box-plots representation of the same dataset.

We used the same experimental paradigm to quantify the same transport parameters for lysosomes as for FNDs. Contrary to our findings for endosome transport, the exposure of neurons to α-synF did not modify lysosome transport parameters (Fig. 3*E*-*H*). However, in cortical neurons exposed to α-synR, lysosome velocity increased by 6% (Fig. 3*E*), and run length increased by 26% (Fig. 3*F*), with no significant change in pausing time (Fig. 3*G*) and a slight increase (by 4%) in pausing frequency (Fig. 3*H*). These results are summarized in Fig. 2-2B.

### Aß assemblies affect the number of cargoes transported along microtubules without significantly modifying trajectory length

We also analyzed the same parameters after exposing DIC20 mouse cortical neurons to either AßF or AßO for 24 hours, and then measuring intracellular transport on DIC21. We used a concentration of 1 μM, which has been reported to have a biological impact (Marshall et al., 2020). The number of FNDs interacting with neurons decreased slightly after exposure to AßF or AßO (by 3% and 7% for AßF and AßO, respectively; Figure 4*A*). This decrease was accompanied by a much larger decrease (56% and 29% for AßF and AßO, respectively) in the fraction of FNDs displaying directed motion (Fig. 4*B*). FND trajectory lengths were almost unaffected by AßF (3% decrease) but were decreased by AßO (Fig. 4*C*, Fig. 4*D*; 13% decrease). We also investigated the effect of Aß assemblies at a lower concentration (0.2 μM), identical to that for α-syn assemblies. Even at this lower concentration, we detected small decreases in the number of FNDs per FoV (Fig. 4-1*A*; 19% and 18% for AßF and AßO, respectively)) and the proportion of FNDs with directed motion (Fig. 4-1*B*; 11% and 18% for AßF and AßO respectively), for both AßF and AßO. Thus, as for α-syn assemblies (Fig. 3), the exposure of cortical neurons to Aß assemblies induces significant major decreases in endosomal transport.

**Figure 4.**
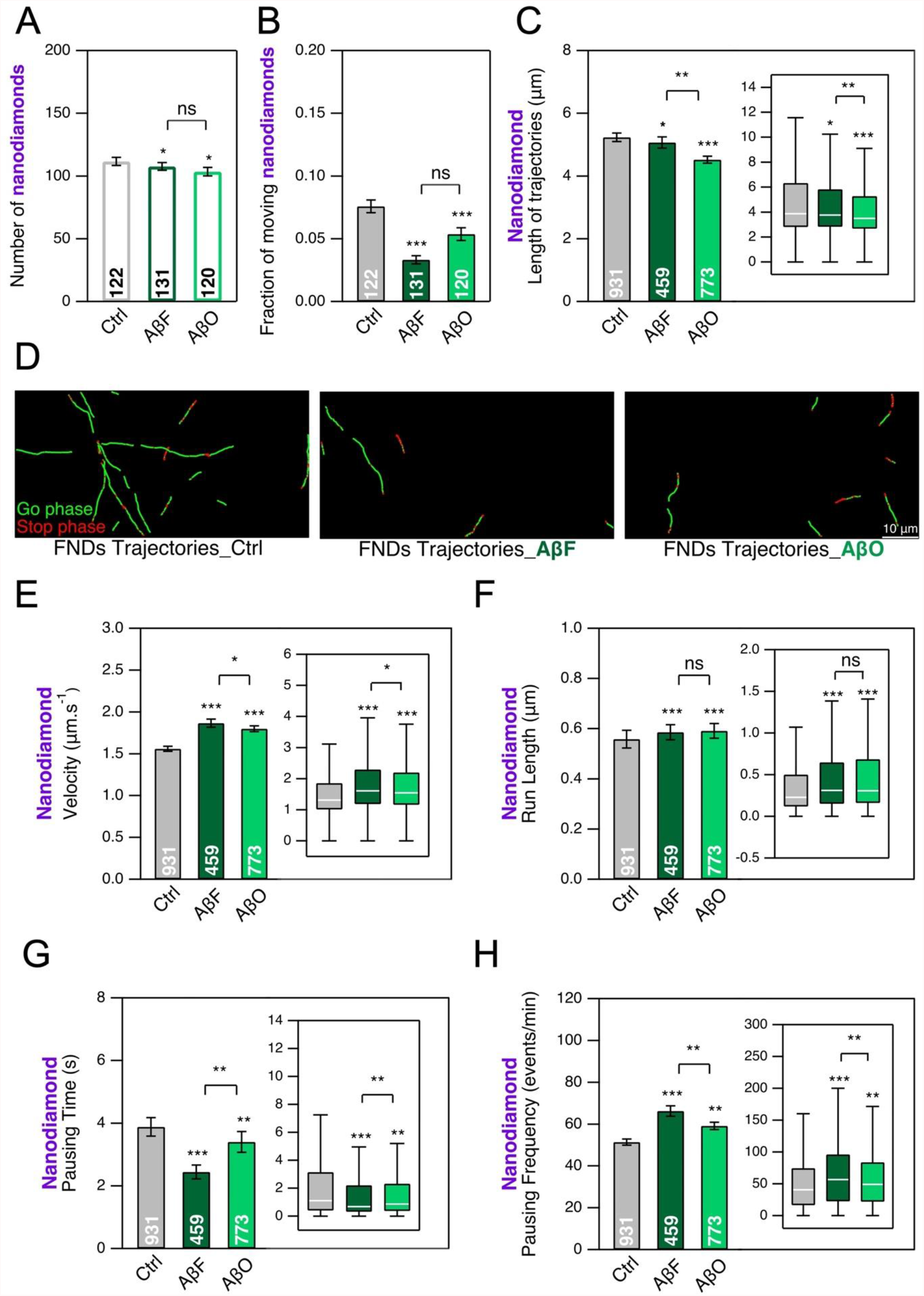
Effect of AßF and AßO on the mobility of endosomes and their transport as measured by tracking FND-containing cargoes in mouse cortical neurons at DIC21. 24 h exposure to AßF and AßO at 1 μM concentration, compared to nothing added control (Ctrl). ***A***, Number of FNDs detected per field-of-view of 40 × 80 μm size during 2 mins of observation. ***B***, Fraction of FNDs-containing cargoes having a directed motion. ***C***, Length of FND trajectories. ***D***, Examples of FND trajectories. Scale bar: 10 μm. ***E***-***H***, Comparison of four transport parameters: curvilinear velocity (***E***), run length (***F***), pausing time (***G***) and pausing frequency (***H***). The number inside the bar represents the total number of FoV (***A, B***) or trajectories (**C, *E***-***H***) analyzed from *n*=6 coverslips (three independent cultures). Inset: box-plots representation of the same dataset. See also Figure 4-1.

We then measured the impact of AßF and AßO on endosomal transport parameters more precisely. We observed increases in FND velocity (Fig. 4*E*; 20% and 15% for AßF and AßO, respectively) and run length (Fig. 4*F*; 5% and 7% for AßF and AßO, respectively), a decrease in pausing time (Fig. 4*G*; 36% and 12% for AßF and AßO, respectively) and an increase in pausing frequency (Fig. 4*H*; 28% and 15% for AßF and AßO, respectively), with more pronounced effects for AßF than for AßO. Interestingly, similar patterns were also observed, for both AßF and AßO, at the lower concentration of 0.2 μM (Fig. 4-1*C*-*F*). Finally, for AßF, the major changes to some transport parameters observed combined to yield only a very slight decrease in trajectory length (Fig 4*C*). The detailed quantitative analysis performed was, therefore, particularly useful. These results are summarized in Fig. 2-2A.

Our investigations of the impact of Aß on lysosomal transport revealed large differences between the two types of assembly: AßF and AßO. The total number of lysosomes detected in a FoV was 50% greater than that for controls in neurons exposed to AßO (Fig. 5*A*), but was similar to that for controls in neurons exposed to AßF. The intracellular transport measurements performed showed that the fraction of moving lysosomes was 1.6-fold lower in neurons exposed to AßF and 5.7-fold lower in neurons exposed to AßO (Fig. 5B; 23% for the control, 14% for AßF and 4.5% for AßO). Lysosome trajectory lengths were shortened only slightly for AßF (5%) and more significantly for AßO (Fig. 5C and Fig. 5D; 17% decrease).

**Figure 5.**
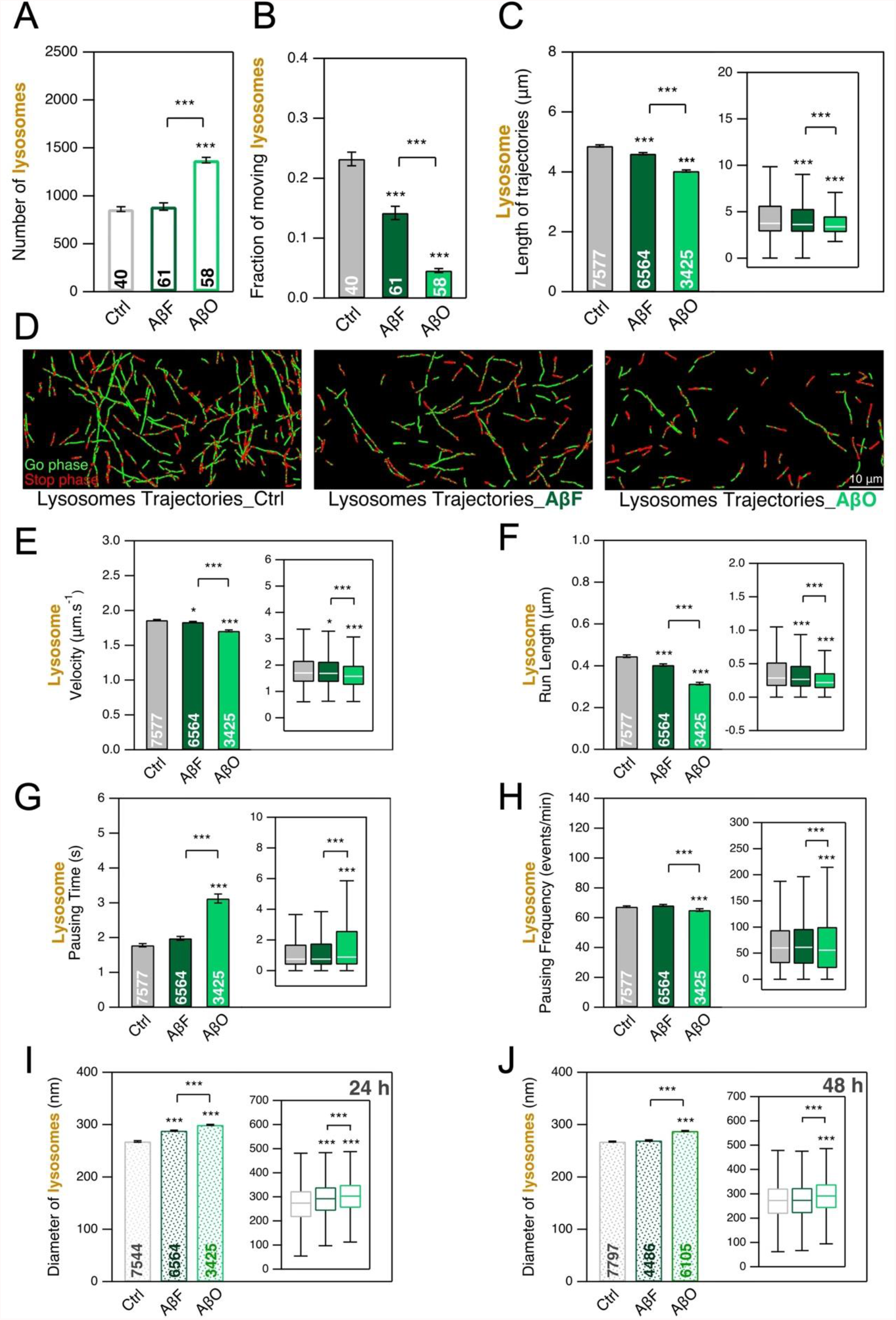
Effect of AßF and AßO on the mobility of LysoTracker-labelled lysosomes and their transport in mouse cortical neurons at DIC21. 24 h exposure to AßF and AßO at 1 μM concentration, compared to nothing added control. ***A***, Number of lysosomes detected per field-of-view of 40 × 80 μm size during 2 mins of observation. ***B***, Fraction of lysosomes having a directed motion. ***C***, Length of lysosome trajectories. ***D***, Examples of lysosome trajectories. Scale bar: 10 μm. ***E****-****H***, Comparison of four transport parameters: curvilinear velocity (***E***), run length (***F***), pausing time (***G***) and pausing frequency (***H***). I-J) Comparison of Lysosome size. The number inside the bar represents the total number of FoV (***A, B***), trajectories (***E***-***H***) and lysosomes (***I, J***) analyzed from *n*=2 coverslips (from one culture). Inset: box-plots representation of the same dataset. See also Figure 5-1, Figure 5-2 and Figure 5-3.

We also measured lysosome transport parameters in the presence of 1 μM AßF or AßO (Fig. 5E-H). For AßF, we observed almost no change in velocity (Fig. 5E), pausing time (Fig. 5G), or pausing frequency (Fig. 5H). By contrast, exposure to AßO led to a 1.7-fold increase in pausing time. Run length decreased significantly for both assemblies (Fig. 5F; 9% and 13% decreases for AßF and AßO, respectively). These results are summarized in Fig. 2-2B.

Changes in lysosome size have been described in the APP mouse transgenic model of Alzheimer’s disease (Gowrishankar et al., 2015). We therefore investigated whether quantifiable changes in lysosome diameter could be detected after 24 or 48 h of exposure to 1 μM AßF or AßO. We detected a slight increase (7%) in lysosome diameter following the exposure of neurons to AßF for 24 h. This increase in size had disappeared by 48 h. By contrast, we observed an increased in lysosome diameter at both time points (11% and 7% at 24 h and 48 h, respectively) in neurons exposed to 1 μM AßO (Fig. 5*I*-*J* and Fig. 5-1*A*-*C*). The finding that AßO triggers an increase in lysosome number and size, and a decrease in lysosome movement is consistent with previous reports (Gowrishankar et al., 2015; Marshall et al., 2020).

Finally, LysoTracker can also label acidic compartments other than lysosomes, including late endosomes, in particular. We therefore repeated the transport experiment with the Magic Red substrate, which reveals cathepsin B protease activity as fluorescence, specifically in lysosomes. For both α-synF (Fig. 5-2) and AßF (Fig. 5-3), the Magic Red-labeled compartments (lysosomes) behaved similarly to the LysoTracker-labeled compartments, for all parameters. We observed, in particular, a decrease in the mobile fraction of ≈40% for both the Magic Red- and LysoTracker-labeled compartments in the presence of the fibrillar assemblies. The colocalization data obtained at 24 h were also similar (*i*.*e*. 4-7% colocalization for α-synF versus 40% for AßF). Thus, the labeling profiles of LysoTracker and Magic Red largely overlapped, and it is reasonable to use LysoTracker puncta density to quantify endocytic activity. These results are summarized in Fig. 2-2C.

### Transport of α-syn and Aß assemblies within cortical neurons

We also assessed the transport of α-syn and Aß assemblies within cortical neurons, while documenting their impact on the dynamics of endosomes and lysosomes. The ATTO 488 dye used to label α-syn and Aß assemblies has no emission spectrum overlap with either FNDs or Lysotracker Deep Red. We were, therefore, able to measure the transport properties of endosomes or lysosomes and the assemblies simultaneously, with two color channels.

We first studied α-syn fibrillar assembly transport (Fig. 6). α-synF and R displayed directed movement, as illustrated by the trajectories in Fig. 6*A*-*B*. We compared these motions to the endosomal transport observed in the presence of α-syn fibrillar polymorphs. The α-syn F and R trajectories were about 29% shorter than those of FNDs (Fig. 6*C*). We also compared the transport parameters and found that velocity (Fig. 6*D*; 5% and 10% for α-synF and α-synR, respectively), run length (Fig. 6*E*; 39% and 46% for α-synF and α-synR, respectively) and pausing time (Fig. 6*F*; 19% and 21% for α-synF and α-synR, respectively) were lower for α-synF- and α-synR-loaded cargoes than for FND-containing endosomes. Conversely, α-synF and α-synR pausing frequencies were greater than those of FNDs (Fig. 6*G*; 19% and 8% for α-synF and α-synR, respectively). The shorter trajectories and run lengths and the longer pausing frequency suggest that cargoes loaded with α-synF- and α-synR are transported less efficiently than those containing FNDs.

**Figure 6.**
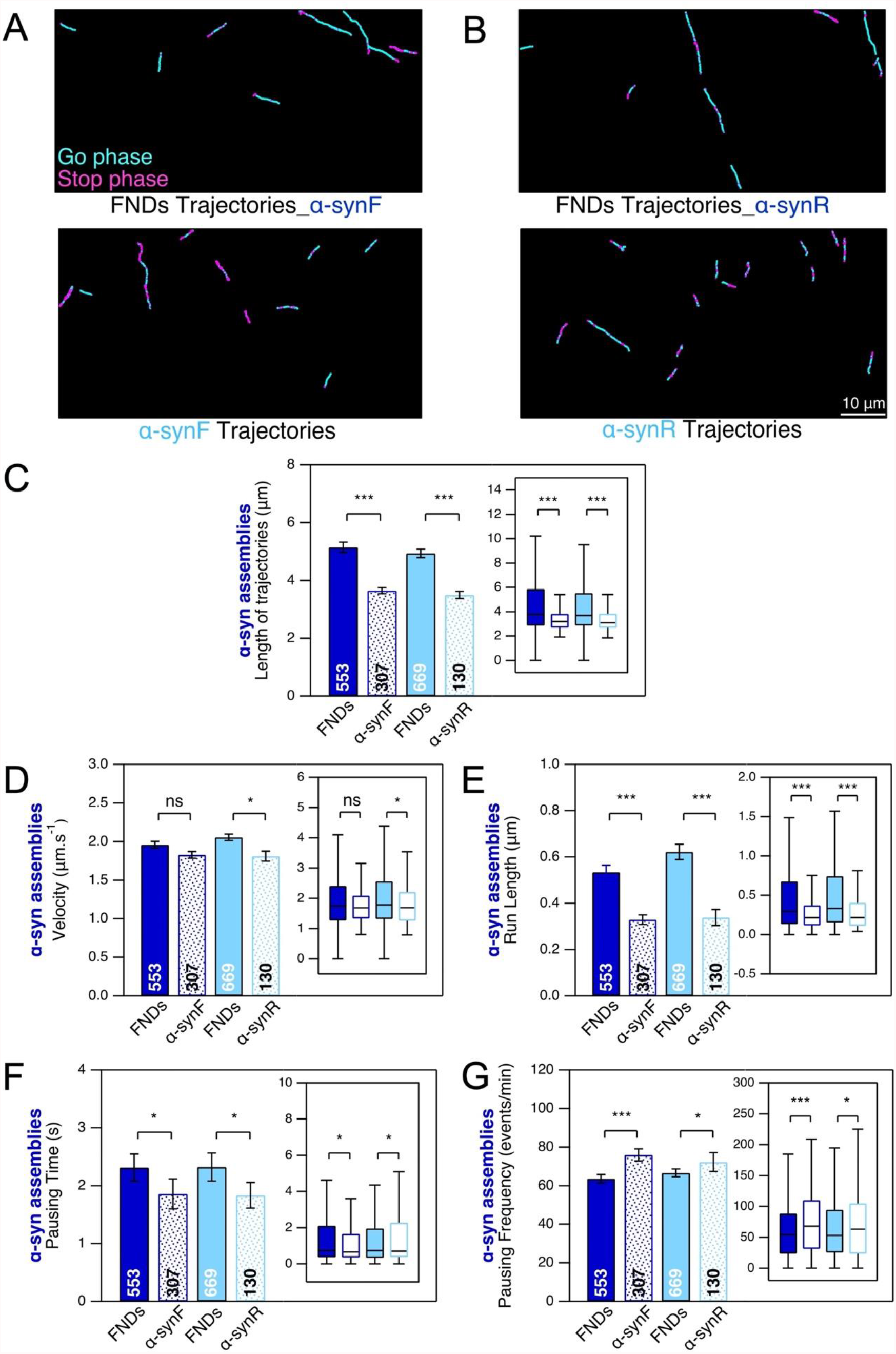
Intraneuronal transport of ATTO 488-labeled α-synF and R in mouse cortical neurons at DIC21. DIC20 cortical neurons were exposed to α-synF and R during 24 h, at concentration of 0.2 μM. ***A***-***B***, Examples of α-synF and R, and FND trajectories (in the presence of α-synF and R). Scale bar: 10 μm. ***C***, Length of α-synF, α-synR and FND trajectories. ***D***-***G***, Comparison of four transport parameters: curvilinear velocity (***D***), run length (***E***), pausing time (***F***) and pausing frequency (***G***). The number inside the bar represents the total number of trajectories analyzed from *n*=8 coverslips (four independent cultures) is indicated in each bar. Inset: box-plots representation of the same dataset.

We also investigated the intraneural transport of ATTO 488-labeled AßF and AßO (Fig. 7) at a concentration of 1 μM, as we detected no fluorescence signal with a concentration of 0.2 μM. Both species displayed directed transport, as shown by the trajectories in Fig. 7*A*-*B*. These trajectories are shorter than those of FNDs in the same conditions (Fig. 7*C*), as for α-syn fibrillar assemblies. AßF and AßO had slightly higher velocities than FNDs (Fig. 7*D*; 8% and 12% for AßF and AßO, respectively), and tended to have a shorter run length (Fig. 7*E*; 7% and 28% for AßF and AßO, respectively). Like α-syn fibrillar assemblies, AßF and AßO had a much shorter pausing time (Fig. 7*F*; 49% and 53% for AßF and AßO, respectively) and a much higher pausing frequency (Fig. 7*G*; 41 and 36% for AßF and AßO, respectively). As for α-syn fibrillar assemblies, these shorter trajectories and run lengths, together with the higher pausing frequency, suggest that cargoes loaded with AßF and AßO are transported less efficiently than those containing FNDs.

**Figure 7.**
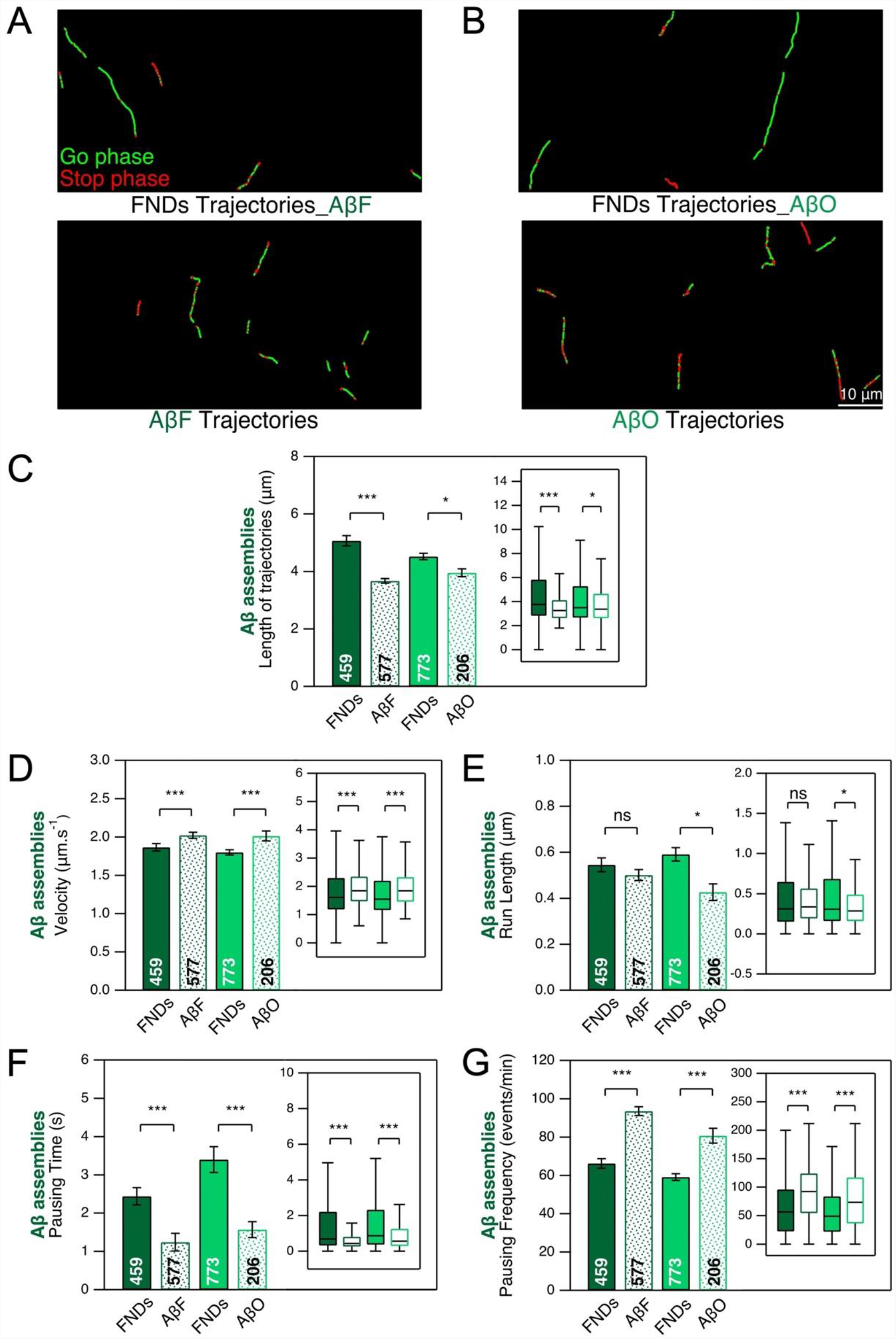
Intraneuronal transport of ATTO 488-labeled AßF and AßO in mouse cortical neurons at DIC21. DIC20 cortical neurons were exposed to AßF and AßO (1 μM) during 24 h. ***A***-***B***, Examples of trajectories. Scale bar: 10 μm. ***C***, Length of AßF, AßO and FND trajectories. ***D***-***G***, Comparison of four transport parameters: curvilinear velocity (***D***), run length (***E***), pausing time (***F***) and pausing frequency (***G***). The number inside the bar represents the total number of trajectories analyzed from *n*=6 coverslips (from three independent cultures) is indicated in each bar. Insets: box plots representation of the same dataset. See also Figure 7-1.

Differences in the uptake of AßO and AßF have recently been reported in cultured neurons (Vadukul et al., 2020). We, therefore, also investigated the related aspect of the number of Aß assembly trajectories per field-of-view at 24 h and 48 h (Fig. 7-1). We observed no differences in trajectory number 24 h after the addition of the assemblies (Fig. 7-1*A*), consistent with Fig. 4B of Vadukul *et al*. (2020). However, at 48 h, there were about 2.5 times as many AßF trajectories as AßO trajectories (Fig. 7-1*B*). These findings contrast with those of Vadukul *et al*. (2020) for the 72 h time point, at which the amount of internalized AßO was about 1.5 times the amount of internalized AßF (sonicated). There are several possible reasons for this discrepancy: (i) we quantified only the moving fraction of Aß assemblies; (ii) we did not consider the 72 h time point, and, finally, (iii) we did not use exactly the same AßF.

Finally, we quantitatively assessed the colocalization of α-syn and Aß assemblies with lysosomes, as a function of neuron exposure time, at 24 and 48 hours (Fig. 8). The proportion of α-syn fibrillar assemblies moving within lysosomes increased from ≈4% at 24 h to 12-14% at 72 h (Fig. 8*A*), but it had already reached ≈41% at 24 h for Aß fibrils, subsequently continuing to increase, up to ≈51% at 48 h (Fig. 8*B*). Colocalization proportions were slightly lower for AßO moving in or with lysosomes, but were much larger than for α-syn fibrillar assemblies. We repeated these analyses with Magic Red labeling in place of LysoTracker for α-synF (Fig. 8-1*A*) and AßF (Fig. 8-1*B*). At 24 h, the results were similar, with ≈40% colocalization of AßF with Magic Red-labeled lysosomes, and only ≈7% colocalization for α-synF.

**Figure 8.**
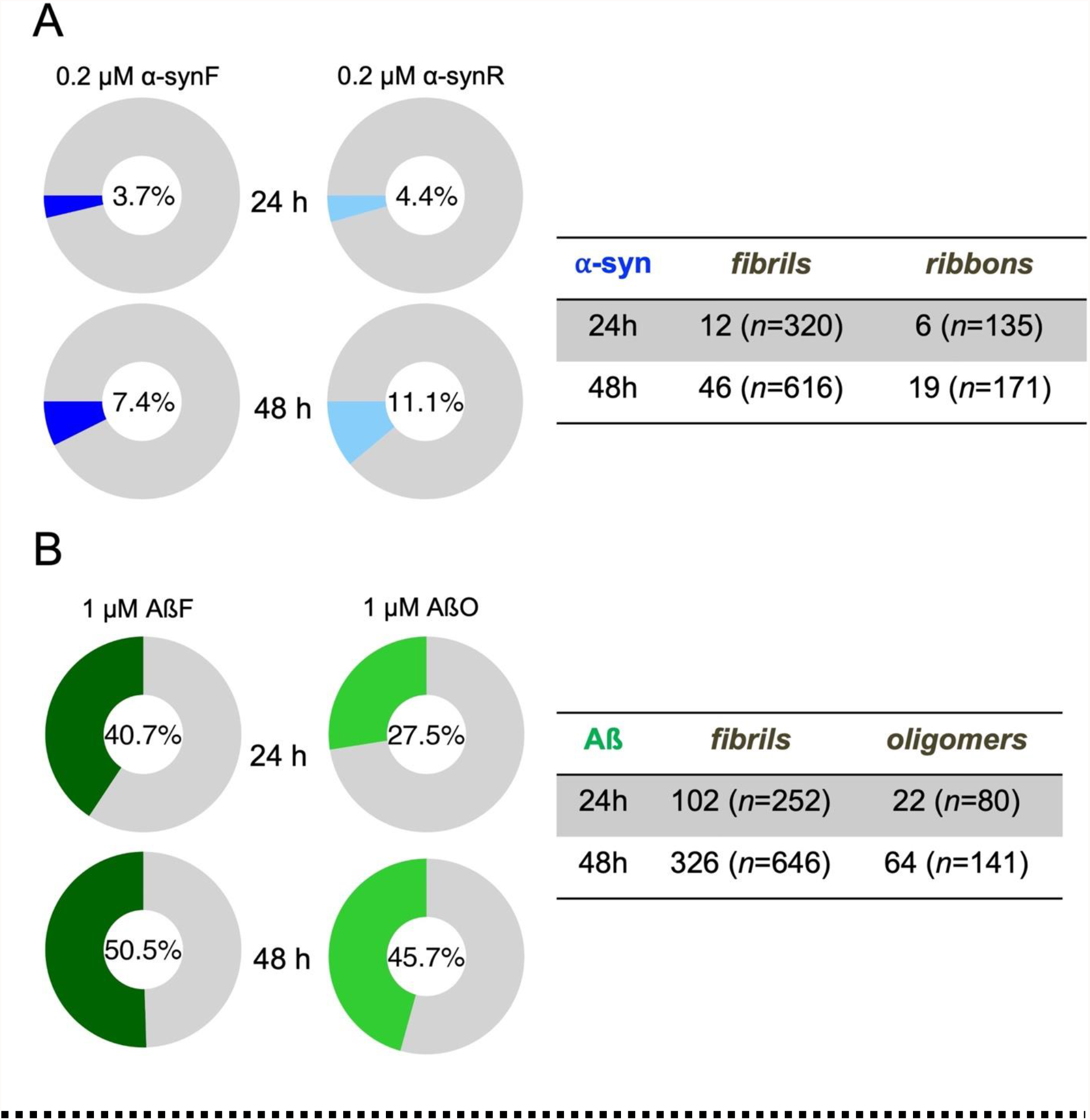
Colocalized events between moving neurodegenerative disease-related molecular species and moving lysosomes at different time points. ***A***, α-synF and α-synR were incubated for 24 and 48 h, at concentration of 0.2 μM. ***B***, AßF and AßO were incubated for 24 and 48 h, at concentration of 1 μM. The number inside the donut plot represents the percentage of moving α-syn or Aß assemblies colocalized with lysosomes (LysoTracker labelled). The table on the right panel indicates the number of neurodegenerative-related molecular species trajectories colocalized with lysosomes. n represents the total number of trajectories. The percentage and number of trajectories in each time point were analyzed from 2 coverslips (from one culture). See also Figure 8-1.

## Discussion

We investigated the generic effects of α-syn fibrillar polymorphs (fibrils and ribbons) and Aß assemblies (oligomers and fibrils) on endosomal and lysosomal movements in mouse cortical neurons. Early endosomes and lysosomes make use of different types of machinery to move in neurons. Furthermore, early endosomes are compartmentalized (dendrites vs. axon) whereas lysosomes are not (Winckler et al., 2018). However, due to the high density of the cultures, it was not possible to identify the compartments (dendrite or axon) in which the tracked vesicles were moving unambiguously. We were, therefore, unable to study the impacts of protein assemblies on axonal and dendritic endolysosomal transport separately.

### Potential consequences of a decrease in the number of cargoes transported at a given time within a cortical neuron

The exposure of cortical neurons to α-syn and Aß assemblies led to large decreases (between 32% and 56%) in the numbers of endosomes and lysosomes moving along neuronal branches (Figs. 1-2). We previously showed that pathogenic α-syn and Aß assemblies bind the plasma membrane, leading to a redistribution of essential membrane proteins (Renner et al. 2010; Shrivastava et al., 2013; Shrivastava et al., 2015). We also reviewed the underlying pathophysiological mechanisms and the mechanisms resulting from pathogenic protein assembly-plasma membrane component interactions (Shrivastava et al., 2017). The decreases observed here may result from changes in membrane dynamics and endocytosis rate.

The exposure of cortical neurons to α-syn and Aß assemblies affected the properties of moving FND-containing endosomes. α-syn fibrillar assemblies increased the velocity of these endosomes by 31-38% and their run length by 80-100% (Fig. 2*E*-*F*), and decreased their pausing time by 40% (Fig. 2*G*), with a smaller increase in pausing frequency (19-25%, Fig. 2*H*). We observed similar, but less marked effects for moving FND-containing endosomes following the exposure of neurons to AßF or AßO (Fig. 4*E*-*H*). These changes reflect an increase in the mobility of moving FND-containing endosomes. Thus, α-synF or α-synR, and AßF or AßO decrease the proportion of moving FND-labeled endosomes whilst increasing the overall mobility of the moving endosomes (Fig. 2-2*A*).

A decrease in the number of moving endosomes or lysosomes can affect protein quality control, limiting the elimination of damaged membrane and cytosolic proteins, protein aggregates, and membranous organelles (Winckler et al., 2018). Furthermoreas lysosomes and late endosomes act as mRNA translation platforms (Cioni et al., 2019; Liao et al., 2019; Fernandopulle et al., 2021), changes in the number of cargoes transported at a given time within a cortical neuron would be expected to have a major effect on the mRNA translation platform in either dendrites or axons. In particular, the regulation of protein synthesis and degradation at the neuronal synapse is local and dynamic, resulting in autonomous modifications to the synaptic proteome during plasticity (Giandomenico et al., 2021). Synaptic function may, therefore be affected if the number of moving lysosomes changes.

### Effect of α-syn and Aß assemblies on lysosome transport

α-synF (Fig. 3E-H) and AßF (Fig. 5E-H) had almost no effect on lysosomal transport parameters, as shown by comparisons with the control. By contrast, α-synR (Fig. 3E-H) and AßO (Fig. 5E-H) induced significant changes in lysosome transport parameters (Fig. 2-2*B*). Furthermore, lysosome size increased significantly in the presence of AßO (Fig. 5I-J). These impairments of lysosomal transport in the context of Alzheimer’s disease are fully consistent with previous reports (Gowrishankar et al., 2015; Marshall et al., 2020). Indeed, using a mouse model of Alzheimer’s disease, (Gowrishankar et al., 2015) provided evidence of axonal lysosome accumulations with locally impaired retrograde axonal transport of lysosome precursors. Similarly, Marshall et al. (2020) found that misfolded Aß_42_ affected the endolysosomal pathway. They reported impairments of the uptake of proteins via a dynamin-dependent endosomal mechanism and the accumulation of lysosomes.

### Intraneuronal transport of neurodegenerative disease-related molecular species

We determined the intraneuronal transport parameters for the pathogenic species α-syn and Aß (Figs. 6-7). The transport of cargoes loaded with α-synF and α-synR was more dynamic than that of cargoes containing FNDs, which had a higher pausing frequency and shorter run-lengths and pausing times (Fig. 6*D*-*G*). AßF- and AßO-containing endosomes had similar characteristics (Fig. 7*D*-*G*), with the addition of an increase in velocity not observed for α-synF and α-synR. These results suggest that similar molecular mechanisms operate in the transport of the two α-syn fibrillar polymorphs, AßF and AßO. However, the seven times larger fraction of AßF and AßO than of α-syn assemblies in moving lysosomes (Fig. 8) also indicates differences in the molecular interactions of Aß assemblies with lysosomes. These results suggest that cellular triage may underlie the differential transport of α-syn and Aß assemblies.

### Potential application in drug discovery assays

We report here the use of a model based on primary mouse cortical neurons, in which we observed a robust decrease in the numbers of vesicles (30-50% decrease; Figs. 2*B*, 3*B*, 4*B* and 5*B*), and lysosomes (five-fold decrease Fig. 5*B*) moving intracellularly in neurons exposed to AßO (Fig. 5*B*). This large decrease probably affected neuron physiology, and the transport of other cargoes, such as mitochondria and RNA granules. This endolysosomal transport impairment endophenotype may be useful for large-scale drug-discovery campaigns (*i*.*e*., >10^5^ compounds), which generally usemore complex human cellular models (Park et al., 2021).

The decrease in cargo transport within cortical neurons described here may result from a direct interaction between pathogenic aggregates and the intraneuronal transport machinery or pathogenic aggregate-mediated changes in the transcription of transport protein genes (Encalada and Goldstein, 2014, Lee et al., 2014; Guo et al., 2020). Analyses of neuronal immunoprecipitates of α-syn fibrillar assemblies and Aß polymorphs may be useful for identifying the molecular partners interacting directly with these protein assemblies. A recent postmortem proteomics study identified proteins for which abundance changed at different stages of Alzheimer’s disease (Li et al., 2021). At early stages, differentially expressed proteins of the “clathrin-coated endocytic vesicle membrane” (GO: 0030669) and the secretory pathway (R-HSA-432720: “Lysosome Vesicle Biogenesis” and R-HSA-432722: “Golgi-Associated Vesicle Biogenesis”) classes were overrepresented. A comparison of the changes in proteome profile between our neuronal model and the results reported by Li et al., 2021 might help to identify druggable targets for increasing the number of cargoes transported.

Finally, we quantified the intraneuronal transport of neurodegenerative disease-associated molecular species (α-syn fibrillar polymorphs, AβF and oligomers), which have transport characteristics different from those of endosomes, for which no molecular characterization is yet available. We found that all these protein assemblies were transported intracellularly in cortical neurons, with very similar quantitative characteristics. We identified no differences in their transport parameters. Further studies will be required to identify a possible common transport mechanism and specific molecules involved in it, with a view to developing ways of selectively inhibiting this mechanism.

These results should also be considered from the standpoint of the prion-like spread of pathogenic protein particles between neurons (Brundin et al., 2010; Hardy and Revesz, 2012; Jucker and Walker, 2018). Selective inhibition may prevent the spread of these neurotoxic species. Advances in the identification of targets involved in the transport of cargoes loaded with pathogenic protein aggregates could potentially lead to the development of novel neuroprotective treatments.

## Acknowledgments

This work was supported by the Joint Program on Neurodegenerative Disease Research and the *Agence National de la Recherche* (TransPathND ANR-17-JPCD-0002 contract awarded to M.S. and R.M.); Program EuroNanoMed3 (MoDiaNo, ANR-18-ENM3-0002 to M.S.). *Fondation pour la Recherche Médicale* (ALZ201912009776 contract awarded to R.M.); Q.L.C. was supported by a PhD scholarship from the Taiwan Ministry for Education and Paris-Saclay University, and the and JPND TransPathND contract (ANR-17-JPCD-0002).

## Extended Figure Legends

**Figure 2-1**. FNDs interaction with neurons and fluorescence intensity of ATTO 488-labeled α-synF with and without (w/o) washing. 24 h exposure of neurons to α-synF at concentration of 0.2 μM, compared to nothing added control. ***A***, Number of FNDs detected per field-of-view of 40 μm x 80 μm during 2 mins of observation for the different conditions. Ctrl: no addition of α-synF; α-synF Wash: addition of α-synF during 24 h and washing of the culture just before addition of FND tracers. Inset: box-plots representation of the same dataset. ***B***, Fraction of FNDs-containing cargoes having a directed motion. The numbers inside the bar in A and B represent the total number of FOV analyzed from *n*=2 coverslips (from one culture). ***C***, Example of FND trajectories in the different conditions. Scale bar: 10 μm. ***D***, Average fluorescence intensity of ATTO 488-labeled α-synF evaluated for 30 μm-long branches, with or w/o washing (*n*= 25 branches from 25 fields-of-view). ***E***, Examples of first frames of TIRF videomicroscopy of ATTO 488-labeled α-synF decoration of DIC21 cortical neurons with (top) or w/o (bottom) washing. Scale bar: 10 μm.

**Figure 2-2**. Summary of the effect of different protein assemblies on the transport parameters of endosomes (***A***), LysoTracker-labeled lysosomes (***B***), and Magic Red-labeled lysosomes (***C***).

**Figure 4-1**. Impact of Aβ fibrils and oligomers at the concentration of 0.2 μM on the number of nanodiamonds interacting with cells (***A***), on the fraction of moving nanodiamonds (***B***) and on the FND-labelled endosome transport parameters (***C*-*F***).

**Figure 5-1**. Lysotracker-labelled compartments size in control (***A***) and 24 h exposure of Aß fibrils (***B***) and oligomers (***C***) at concentration of 1 μM. Scale bar: 2 μm.

**Figure 5-2**. Effect of α-synF on the mobility of Magic Red-labelled lysosomes and their transport in mouse cortical neurons at DIC21, after 24 h exposure to α-synF at concentration of 0.2 μM, compared to nothing added control. ***A***, Number of lysosomes detected per field-of-view of 40 μm x 80 μm size during 2 mins of observation. ***B***, Fraction of lysosomes having a directed motion. ***C***, Trajectory length. ***D***, Examples of lysosome trajectories. Scale bar: 10 μm. ***E-H***, Comparison of four transport parameters: **E)** curvilinear velocity (***E***), run length (***F***), pausing time (***G***) and pausing frequency (***H***). ***I***, Diameter of Magic Red-labelled lysosomes. General remarks: the number inside the bar represents the total number of FoV (***A, B***), trajectories (***E***-***I***). Inset: box-plots representations of the same datasets (***C***-***I***).

**Figure 5-3**. Effect of AßF on the mobility of Magic Red-labelled lysosomes and their transport in mouse cortical neurons at DIC21 after 24 h exposure to AßF at concentration of 1 μM, compared to nothing added control. ***A***, Number of lysosomes detected per field-of-view of 40 μm x 80 μm size during 2 mins of observation. ***B***, Fraction of lysosomes having a directed motion. ***C***, Trajectory length. ***D***, Examples of lysosome trajectories. Scale bar: 10 μm. (***E***-***H***) Comparison of four transport parameters: curvilinear velocity (***E***), run length (***F***), pausing time (***G***) and pausing frequency (***H***). ***I***, Comparison of Magic Red-labelled lysosome size. General remarks: The number inside the bar represents the total number of FoV (***A, B***) and trajectories (***E***-***I***) Inset: box-plots representation of the same dataset.

**Figure 7-1**. Number of AßF and AßO trajectories detected per field-of-view of 40 × 80 μm during 2 mins of observation. Cortical neurons were exposure to AßF and AßO (1μM) during 24 h (***A***) and 48 h (***B***). The number inside the bar represents the total number of FOV analyzed from *n*= 6 coverslips (from three cultures) for 24 h and *n*=2 coverslips (from one culture).

**Figure 8-1**. Fraction of protein assemblies, α-synF (***A***) or AßF (***B***) trajectories colocalizing with Magic Red puncta trajectories, as inferred from the tables indicating the number of protein assembly trajectories colocalizing with Magic Red puncta and the total number of assembly trajectories in parenthesis. Data from 2 coverslips and one culture in both α-synF and Aß cases.

## References

Alam P, Bousset L, Melki R, Otzen DE (2019) α-synuclein oligomers and fibrils: a spectrum of species, a spectrum of toxicities. J Neurochem 150(5):522–534.

Brahic M, Bousset L, Bieri G, Melki R, Gitler AD (2016) Axonal transport and secretion of fibrillar forms of α-synuclein, Aβ<sub>42 </sub>peptide and HTTExon 1. Acta Neuropathol 131(4):539–48.

Braak H, Braak E (1991) Neuropathological stageing of Alzheimer-related changes. Acta Neuropathol 82:239–259.

Braak H, Del Tredici K, Ru b U, de Vos RAI, Jansen Steur ENH, Braak E (2003) Staging of brain pathology related to sporadic Parkinson’s disease. Neurobiol Aging 24:197–21.

Brundin P, Melki R, Kopito R (2010) Prion-like transmission of protein aggregates in neurodegenerative diseases. Nat Rev Mol Cell Biol 11(4):301–7.

Cioni JM, Lin JQ, Holtermann AV, Koppers M, Jakobs MAH, Azizi A, Turner-Bridger B, Shigeoka T, Franze K, Harris WA, Holt CE (2019) Late Endosomes Act as mRNA Translation Platforms and Sustain Mitochondria in Axons. Cell 176(1-2):56–72.e15.

Crocker JC, Grier DG (1996) Methods of Digital Video Microscopy for Colloidal Studies. J Colloid Interf Sci 179:298–310.

Encalada SE, Goldstein LS (2014) Biophysical challenges to axonal transport: motor-cargo deficiencies and neurodegeneration. Annu Rev Biophys 43:141–69

Fernandopulle MS, Lippincott-Schwartz J, Ward ME (2021) RNA transport and local translation in neurodevelopmental and neurodegenerative disease. Nat Neurosci 24(5):622–632.

Ghee M, Melki R, Michot N, Mallet J (2005) PA700, the regulatory complex of the 26S proteasome, interferes with alpha-synuclein assembly. FEBS J 272:4023–4033.

Giandomenico SL, Alvarez-Castelao B, Schuman EM (2021) Proteostatic regulation in neuronal compartments. Trends Neurosci 3:S0166–2236(21)00161-2.

Golde TE, Borchelt DR, Giasson BI & Lewis J (2013) Thinking laterally about neurodegenerative proteinopathies. J. Clin. Invest 123:1847–1855.

Gowrishankar S, Yuan P, Wu Y, Schrag M, Paradise S, Grutzendler J, De Camilli P, Ferguson SM (2015) Massive accumulation of luminal protease-deficient axonal lysosomes at Alzheimer’s disease amyloid plaques. Proc Natl Acad Sci U S A 14;112(28):E3699–708.

Guo W, Stoklund Dittlau K, Van Den Bosch L (2020) Axonal transport defects and neurodegeneration: Molecular mechanisms and therapeutic implications. Semin Cell Dev Biol 99:133–150.

Hardy J, Revesz T (2012) The spread of neurodegenerative disease. N Engl J Med 366(22):2126–8.

Haziza S, Mohan N, Loe-Mie Y, Lepagnol-Bestel AM, Massou S, Adam MP, L. XL, Viard J, Plancon C, Daudin R, Koebel P, Dorard E, Rose C, Hsieh FJ, Wu CC, Potier B, Herault Y, Sala C, Corvin A, Allinquant B, Chang HC, Treussart F, Simonneau M (2017) Fluorescent nanodiamond tracking reveals intraneuronal transport abnormalities induced by brain-disease-related genetic risk factors. Nat Nanotechnol 12(4):322–328.

Jucker M, Walker LC (2018) Propagation and spread of pathogenic protein assemblies in neurodegenerative diseases. Nat Neurosci 21(10):1341–1349.

Lee WC, Yoshihara M, Littleton JT (2004) Cytoplasmic aggregates trap polyglutamine-containing proteins and block axonal transport in a Drosophila model of Huntington’s disease. Proc Natl Acad Sci U S A 101(9):3224–9.

Li X, Tsolis KC, Koper MJ, Ronisz A, Ospitalieri S, von Arnim CAF, Vandenberghe R, Tousseyn T, Scheuerle A, Economou A, Carpentier S, Otto M, Thal DR (2021) Sequence of proteome profiles in preclinical and symptomatic Alzheimer’s disease. Alzheimers Dement.

Liao YC, Fernandopulle MS, Wang G, Choi H, Hao L, Drerup CM, Patel R, Qamar S, Nixon-Abell J, Shen Y, Meadows W, Vendruscolo M, Knowles TPJ, Nelson M, Czekalska MA, Musteikyte G, Gachechiladze MA, Stephens CA, Pasolli HA, Forrest LR, St George-Hyslop P, Lippincott-Schwartz J, Ward ME (2019) RNA Granules Hitchhike on Lysosomes for Long-Distance Transport, Using Annexin A11 as a Molecular Tether. Cell 19;179(1):147–164.e20.

Marshall KE, Vadukul DM, Staras K, Serpell LC (2020) Misfolded amyloid-beta-42 impairs the endosomal-lysosomal pathway. Cell Mol Life Sci 77(23):5031–5043.

De Matteis MA, Luini A (2011) Mendelian disorders of membrane trafficking. N Engl J Med 8;365(10):927–38.

Millecamps S, Julien JP (2013) Axonal transport deficits and neurodegenerative diseases. Nat Rev Neurosci 14(3):161–76.

Morfini GA, Burns M, Binder LI, Kanaan NM, LaPointe N, Bosco DA, Brown RH, Brown H, Tiwari A, Hayward L, Edgar J, Nave K-A, Garberrn J, Atagi Y, Song Y, Pigino G, Brady ST (2009) Axonal Transport Defects in Neurodegenerative Diseases. J Neurosci 29:12776–12786.

Park J-C, Jang S-Y, Lee D, Lee J, Kang U, Chang H, Kim HJ, Han S-H, Seo J, Choi M, Lee DY, Byun MS, Yi D, Cho K-H, Mook-Jung I (2021) A logical network-based drug-screening platform for Alzheimer’s disease representing pathological features of human brain organoids. Nat Commun 12:280.

Peelaerts W, Bousset L, Van der Perren A, Moskalyuk A, Pulizzi R, Giugliano M, Van den Haute C, Melki R, Baekelandt V (2015) α-Synuclein strains cause distinct synucleinopathies after local and systemic administration. Nature 522(7556):340–4.

Renner M, Lacor PN, Velasco PT, Xu J, Contractor A, Klein WL, and Triller A (2010) Deleterious effects of amyloid b oligomers acting as an extracellular scaffold for mGluR5. Neuron 66:739–754.

Saez-Atienzar S, Masliah E. Cellular senescence and Alzheimer disease: the egg and the chicken scenario. Nat Rev Neurosci 21(8):433–444.

Sardana R, Emr SD (2021) Membrane Protein Quality Control Mechanisms in the Endo-Lysosome System. Trends Cell Biol 31:269–283.

Saudou F, Humbert S (2016) The Biology of Huntingtin. Neuron 89(5):910–926.

Schindelin J, Arganda-Carreras I, Frise E, Kaynig V, Longair M, Pietzsch T, Preibisch S, Rueden C, Saalfeld S, Schmid B, Tinevez J-Y, White DJ, Hartenstein V, Eliceiri K, Tomancak P, Cardona A (2012) Fiji: an open-source platform for biological-image analysis. Nat Methods 9:676–682.

Shrivastava AN, Aperia A, Melki R, Triller A. (2017) Physico-Pathologic Mechanisms Involved in Neurodegeneration: Misfolded Protein-Plasma Membrane Interactions. Neuron 95(1):33–50.

Shrivastava AN, Bousset L, Renner M, Redeker V, Savistchenko J, Triller A, Melki R. (2020) Differential Membrane Binding and Seeding of Distinct α-Synuclein Fibrillar Polymorphs. Biophys J 118(6):1301–1320.

Shrivastava AN, Kowalewski J.M., Renner M., Bousset L, Koulakoff A, Melki R, Giaume C, and Triller A (2013) ß-amyloid and ATP-induced diffusional trapping of astrocyte and neuronal metabotropic glutamate type-5 receptors. Glia 61:1673–1686.

Shrivastava AN, Redeker V, Fritz N, Pieri L, Almeida LG, Spolidoro M, Liebmann T, Bousset L, Renner M, Léna C, Aperia A, Melki R, Triller A. (2015) alpha-synuclein assemblies sequester neuronal alpha3-Na+/K+-ATPase and impair Na+ gradient. EMBO J 34(19):2408–23.

Shrivastava AN, Redeker V, Pieri L, Bousset L, Renner M, Madiona K, Mailhes-Hamon C, Coens A, Buée L, Hantraye P, Triller A, Melki R. (2019) Clustering of Tau fibrils impairs the synaptic composition of α3-Na+/K+-ATPase and AMPA receptors. EMBO J 38(3):e99871.

Soto C, Pritzkow S (2018) Protein misfolding, aggregation, and conformational strains in neurodegenerative diseases. Nat. Neurosci 21:1332–1340.

Stokin GB, Lillo C, Falzone TL, Brusch RG, Rockenstein E, Mount SL, Raman R, Davies P, Masliah E, Williams DS, Goldstein, LS. (2005) Axonopathy and transport deficits early in the pathogenesis of Alzheimer’s disease. Science 307:1282–1288.

Trackpy (2019), doi: 10.5281/zenodo.3492186

Valm AM, Cohen S, Legant WR, Melunis J, Hershberg U, Wait E, Cohen AR, Davidson MW, Betzig E, Lippincott-Schwartz J. (2017) Applying systems-level spectral imaging and analysis to reveal the organelle interactome. Nature 546(7656):162–167.

Victoria GS, Zurzolo C. (2017) The spread of prion-like proteins by lysosomes and tunneling nanotubes: Implications for neurodegenerative diseases. J Cell Biol 216(9):2633–2644.

Volpicelli-Daley LA, Gamble KL, Schultheiss CE, Riddle DM, West AB, Lee VM. (2014) Parkinson’s disease Formation of α-synuclein Lewy neurite-like aggregates in axons impedes the transport of distinct endosomes. Mol Biol Cell 25(25):4010–23.

Walker LC, Jucker M. (2015) Neurodegenerative diseases: expanding the prion concept. Annu Rev Neurosci 38:87–103.

Walsh DM, Thulin E, Minogue AM, Gustavsson N, Pang E, Teplow DB, Linse S (2009) A facile method for expression and purification of the Alzheimer’s disease-associated amyloid beta-peptide. FEBS J 276:1266–1281.

Winckler B, Faundez V, Maday S, Cai Q, Almeida CG, Zhang H (2018) The Endolysosomal System and Proteostasis: From Development to Degeneration. J Neurosci 38:9364–9374.

